# Reservoir Computing with Ultra-Sparse Rings

**DOI:** 10.64898/2026.02.21.707238

**Authors:** Afroditi Talidou, Wilten Nicola

## Abstract

Many existing models of computation in recurrent neural networks assume dense, unconstrained initial connectivity, where any pair of neurons may be coupled to generate the rich dynamics needed for learning complex temporal patterns. Inspired by invertebrate circuits that often exhibit ring-like connectivity, we show that computation can occur in ultra-sparse spiking and rate reservoirs that are initially coupled as simple unidirectional rings. In contrast to standard recurrent networks, the total number of network parameters in these ring networks scales only linearly with network size, while still producing rich feature sets. We demonstrate that such networks can successfully reproduce a range of dynamical systems tasks, including oscillations, multi-stable switches, and low-dimensional chaotic attractors. Our findings show that structured spatio-temporal dynamics naturally arising from large ring topologies, often observed in invertebrate circuits, are a sufficient mechanism for learning different types of attractors.

## Introduction

A key feature of biological intelligence in both vertebrate and invertebrate systems is the ability to learn, organize, and reliably generate complex sequences. Learned behaviors such as language acquisition and skilled motor performance rely on neural systems that generate highly structured activity over time. This capability emerges from the collective dynamics of interacting neurons whose ongoing activity enables the integration of information over time and space to shape precise behavioral outputs. Recurrent neural network models offer a powerful abstraction for studying and replicating these temporally dependent processes due to their inherent dynamical properties [1–11].

Recurrent networks are frequently formulated with one key assumption: unconstrained structural connectivity between neurons, that is, any two neurons in a network have some probability of sharing a connection. When employed in reservoir computing, a network consisting of *N* neurons and possessing this unconstrained connectivity requires *O*(*N*^2^) parameters, or for vanishingly sparse networks, 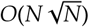 parameters, whether trained or not. The connectivity is also often balanced between positive and negative weights, which produces a dynamically rich state with irregular fluctuations. These irregular and asynchronous dynamics are mathematically characterized as hyperchaotic, with some of the key predictions of excitatory-inhibitory (E-I) balanced computational models having been validated in both *in vitro* [12] and *in vivo* [13] experiments. Such dynamics have been proposed to provide cortical networks with a rich, high-dimensional substrate for computation. This view has strongly influenced training strategies for recurrent networks, including reservoir-based approaches which use the intrinsic variability of strongly recurrent systems to learn and generate complex temporal patterns [14–28]. In these frameworks, effective learning is typically associated with “taming the chaos” that emerges from balanced connectivity.

However, while mammalian circuits may be structurally less constrained, invertebrate circuits often have considerable anatomical constraints. For example, head direction circuits in zebrafish and Drosophila are organized as circular or ring-like networks [29–32] that give rise to continuous attractor dynamics. Indeed, anatomical ring structures are observed across multiple invertebrate species such as C. Elegans [33], jellyfish [34], and starfish [35]. Further, computational studies have shown that various forms of ring coupling can lead to rich asynchronous dynamics that may be utilizable for reservoir computing [37–41]. Given the ubiquity of ring-like structures in those circuits, how are these organisms able to produce complex behaviours with such constrained connectivity?

Inspired by invertebrate models of computation, we show that recurrent neural networks with ring topologies can be reliably trained on a variety of tasks using reservoir computing, performing in many cases as well as classical reservoirs. We trained both networks of spiking and rate neurons organized as ring architectures with the First Order Reduced and Controlled Error (FORCE) algorithm to reproduce complex target dynamics. Instead of requiring *N*^2^ connection weights to drive the initial reservoir, sufficiently rich dynamics emerge with only *O*(*N*) connections constrained to ring-like structures. When trained, the entire system utilizes only *O*(*N*) parameters. Further, we found that inhibition must dominate in spiking networks arranged as rings, since rings with excitatory dominance were unstable. Despite the absence of dense E-I balance, these networks reliably learn and generate rich dynamics, demonstrating that dense connectivity is not a prerequisite for learning in biological and artificial systems, with asynchronous/irregular dynamics reliably emerging from rings. Our results establish a framework for applying supervised learning to recurrent networks in ultra-sparsely connected neural networks, and provide *in silico* evidence for why ring structures may be so ubiquitous in invertebrate circuits: rings produce rich enough dynamics for computation with a fraction of the metabolic requirements. The use of rings instead of dense connectivity has considerable impact for hardware instantiations of reservoir computing, where the computational constraints of neuromorphic hardware may necessitate more sparsely connected reservoirs.

## Results

### Performance of the FORCE method in training rate networks

We first consider networks of rate neurons. In such networks trained with reservoir computing [1], the synaptic weight matrix *ω*, can be expressed as the sum of static and learned components:

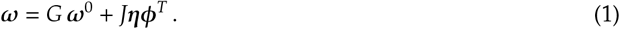

The static component is defined by the static weight matrix *ω*^0^, which is a random matrix scaled by the gain parameter *G*, while the learned component is given by the product of a static random encoder *η* and a learned decoder *ϕ*, scaled by *J*. Classically, the coupling parameter *G* should be scaled to initialize networks near the *edge of chaos* for efficient and rapid learning as the dynamics of neurons in this state are uncorrelated with each other, and rich enough to produce the dynamical features required for learning different attractor states [43–45]. The decoders *ϕ* are learned online using the FORCE algorithm which uses Recursive Least Squares (RLS) to update the decoders [46], while the encoders *η* determine each neuron’s coupling to the learned feedback (see Methods). Critically, the learned component *ηϕ*^*T*^ only contains *O*(*N*) parameters, as it is a low-rank matrix, while *ω*^0^ contains *O*(*N*^2^) parameters, or if the neurons are connected with a probability 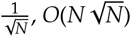 parameters.

Here, we considered rate networks with

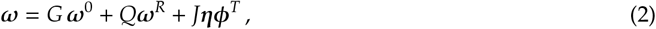

where *ω*^*R*^ has a ring structure (see Methods). When *G* = 0, the system displays a pure ring structure with a low-rank feedback term, and only has *O*(*N*) parameters, while when *Q* = 0, the system is a classical E-I balanced reservoir with *O*(*N*^2^) parameters. Further, the eigenvalues of *ω*^0^ uniformly occupy a circular region in the complex plane of radius *ρ*, and optimal learning occurs near *ρ* ≈ 1, that is, near the edge of chaos. On the other hand, the eigenvalues of *ω*^*R*^ occupy only the “shell” of that circular region, with radius *ρ*. In this work, we show that (i) only the eigenvalue shell has to lie near the edge of chaos, to produce asynchronous dynamics, with the eigenvalue shell generated by ring-connectivity, and (ii) the resulting dynamics, whether chaotic or not, can be tamed to compute any dynamical system.

First, we explored a range of values for *G* and *Q* to identify parameter regimes in which FORCE achieves low error for either ring (*G* = 0), or dense (*G* ≠ 0) configurations (Fig. 1a). Networks with varying values of *G* and *Q* were trained to mimic the FitzHugh-Nagumo oscillator (see Methods). Performance, quantified by the root mean squared error (RMSE), was optimal for values with 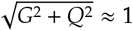, near the edge of chaos. Outside this regime, the RMSE increased substantially. Notably, low error was also obtained in networks that were initially coupled as pure rings, that is, when *G* = 0. This observation implies that the *O*(*N*^2^) parameters and the dense, uniform distribution of eigenvalues within a circle of radius 1 can be substituted with *O*(*N*) parameters of the ring network and a shell of eigenvalues near the onset of instability (*ρ* ≈ 1).

**Figure 1:**
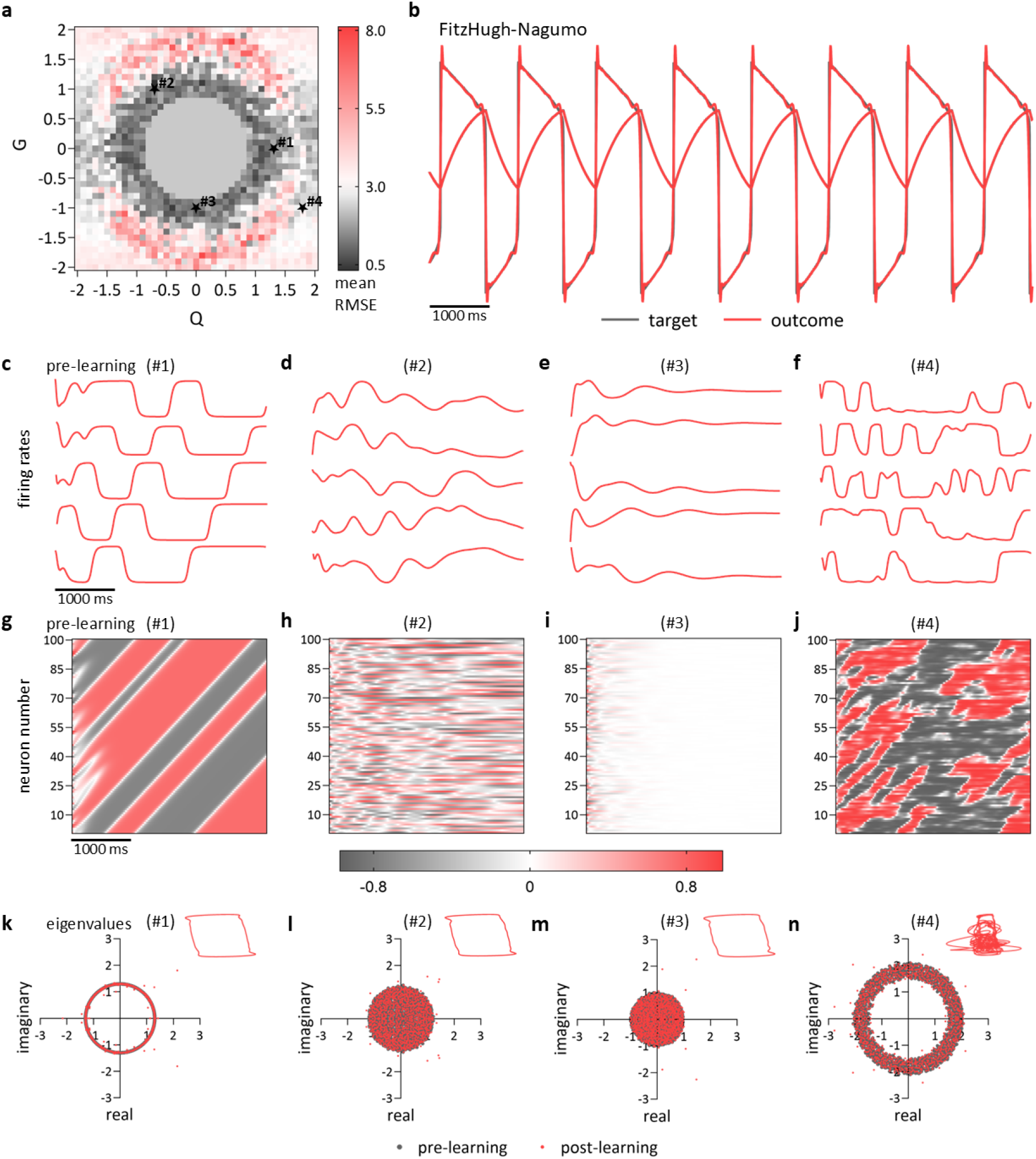
Rate network performance. **a**. In the (*Q, G*) parameter space, networks of *N* = 1000 neurons were trained to reproduce the dynamics of the FitzHugh-Nagumo system. The heatmap represents the mean value of the root mean square errors (RMSE) of the trained networks. The stars indicate the points that were picked for further study: Point #1 = (1.3, 0), Point #2 = (−0.7, 1), Point #3 = (0, −1) and Point #4 = (1.8, −1). **b**. Time series of the two components of the FitzHugh-Nagumo system post-learning. The grey line is the target while the red line is the network outcome. **c-f**. Firing rates of the first five neurons pre-learning for each of the four points picked above. **g-j**. Corresponding heatmaps of the firing rates for the first 100 neurons pre-learning. **k-n**. Spectrum of the eigenvalues for the effective weight matrix for each of the four points. The grey dots represent eigenvalues pre-learning and the red ones post-learning. The insets at the top-right corners are the phase portraits of the FitzHugh-Nagumo system.

To examine this result in further detail, we selected four pairs of *Q* and *G* from distinct regions of the (*Q, G*) parameter space. For Point #1, (*Q, G*) = (1.3, 0), the network dynamics closely match the target dynamics (Fig. 1b) in a ring-like configuration, where the initial firing rates exhibit structured spatiotemporal patterns, which are sufficient to form a functional reservoir that enables learning of the target signal (Fig. 1c,g). The spectrum forms a circle of *ρ* ≈ 1 with eigenvalues occupying only the shell both before and after training. Following training, most eigenvalues remain distributed along this shell, although there are outliers (Fig. 1k).

Next, we considered regimes that were hybrids of rings/dense balanced connectivity. For example, in Point #2, (*Q, G*) = (−0.7, 1), learning exploits the presence of rich chaotic dynamics. Prior to training, firing rates are heterogeneous and time-varying (Fig. 1d,h). After training, the firing rates become coherent (Supplementary Fig. S1b), and the network successfully reproduces the FitzHugh-Nagumo dynamics, as shown in the inset of Fig. 1l and Supplementary Fig. S1m. Following training, the eigenvalues are distributed within a circle of radius one (Fig. 1l), with outliers lying outside of this circle.

Point #3, (*Q, G*) = (0, −1), corresponds to the case in which the ring structure is absent. In this regime, firing rates exhibit transient chaotic activity prior to training, after which they decay to 0 (Fig. 1e,i). Note that as *N* → ∞, the transition to self-sustained chaotic fluctuations occurs at *G* = ±1 (through symmetries induced by using the tanh transfer function). Nevertheless, this transient activity appears sufficient to establish a functional reservoir. When RLS is turned on, firing rates become coherent and remain so after RLS is turned off (Supplementary Fig. S1c). The network reproduces the target dynamics with only a small error (Supplementary Fig. S1n). Following learning, the eigenvalue spectrum forms a circle of radius 1, with a small number of leading eigenvalues lying outside the circle in the complex plane (Fig. 1m). Notably, for points #1, #2, and #3, the post-training spectra exhibit similar qualitative features, namely the presence of outlier eigenvalues that modify the network dynamics. These outliers arise from low-dimensional perturbations [47, 48].

Point #4, (*Q, G*) = (1.8, −1), lies outside the parameter regime in which FORCE performs well. In this case, firing rates exhibit strongly chaotic spatio-temporal dynamics prior to training (Fig. 1f,j), and the network fails to reproduce the target signal (Supplementary Fig. S1o). The corresponding eigenvalue spectrum forms a torus with an outer radius of approximately 2 (Fig. 1n). Firing rate patterns pre- and post-learning for each of the four points are shown in Supplementary Fig. S1a-d, and Fig. S1i-l, along with rate phase portraits (Supplementary Fig. S1e-h).

As shown above, FORCE enables a rate network to reproduce a simple oscillator with an initial ring network as the “reservoir” (see point #1). We next sought to determine if ring-rate networks could learn more complex dynamics by training a network to reproduce the Lorenz attractor (Fig. 2). Here, the initial reservoir was coupled as a ring (Fig. 2a) with RLS used to learn the decoders *ϕ* (Fig. 2b-e). RLS is activated following a brief initialization period and remains active throughout training, after which it is deactivated. Upon deactivation, the decoders remain fixed at their last updated values (Fig. 2b). To elucidate how the network acquires these dynamics, we computed the eigenvalue spectrum of the weight matrix after learning (Fig. 2c). The resulting spectrum after learning exhibits a fuzzy circle of eigenvalues near the original shell. Prior to learning, network activity is highly structured and asynchronous, whereas it becomes coherent following training (Fig. 2d). During training, the network dynamics closely track the target Lorenz dynamics. However, once RLS is deactivated, the network output diverges from the target trajectory due to the chaotic nature of the Lorenz system (Fig. 2e). Nevertheless, the trajectories produced by the trained network remain similar to the original Lorenz trajectories (Fig. 2f). To further confirm that the model learned the Lorenz attractor, we compared the return map constructed from successive maxima of the Lorenz system and found that the trained network reproduces the Lorenz return map (Fig. 2g), indicating that a ring-rate network can learn to perform chaotic forecasting. Therefore, we showed that reservoirs with ring architectures can readily learn complex dynamics.

**Figure 2:**
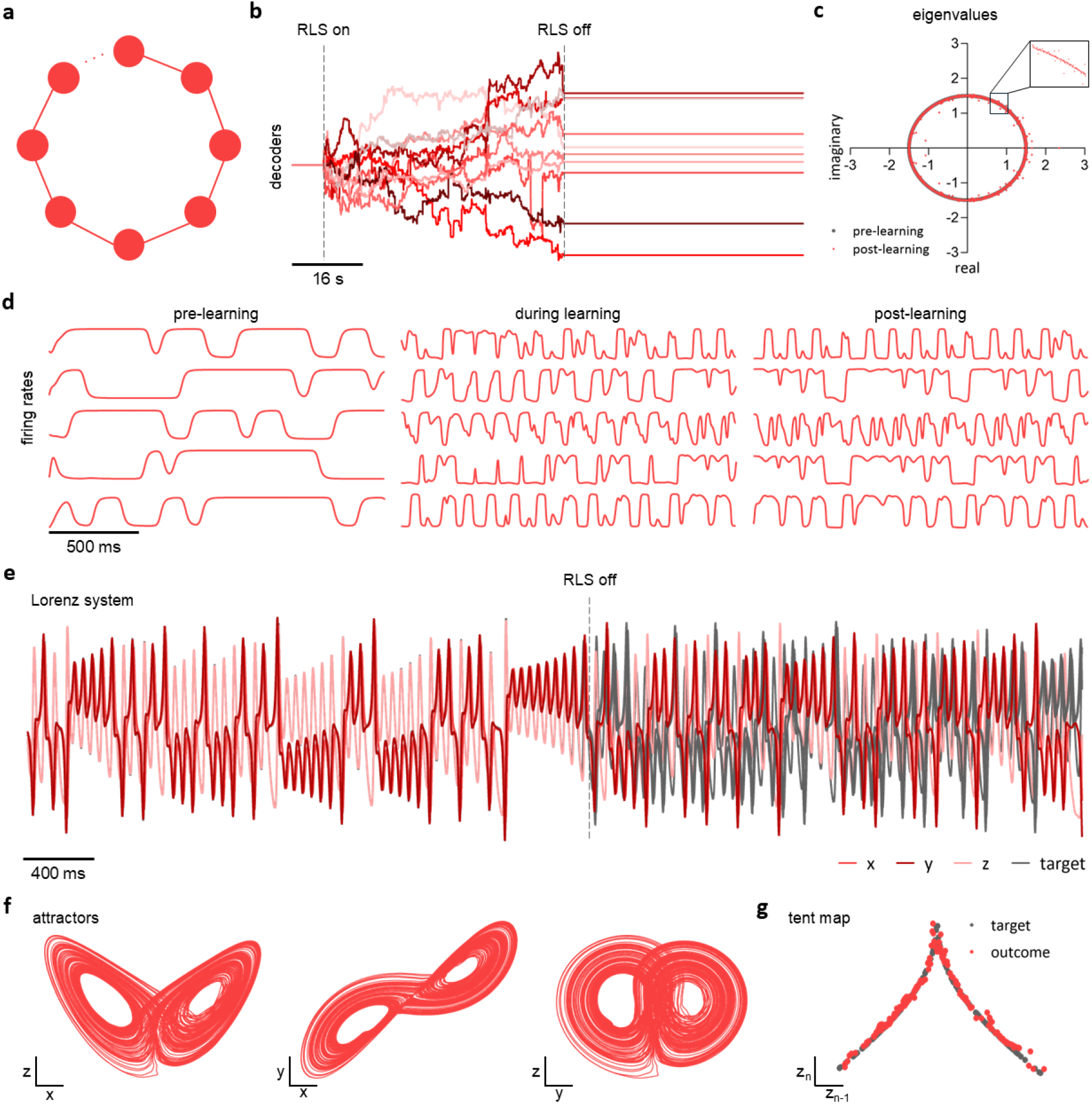
Simple ring rate networks learn chaotic dynamics. **a**. Schematic representation of the complete ring network. The network consists of *N* = 2000 neurons recursively connected. **b**. Decoders of 10 randomly selected neurons, before, during and after FORCE training. The training period lasts for 7.5 seconds. **c**. The eigenvalues are located along the “shell” of a circular domain. Grey dots represent the eigenvalues before training and red dots after FORCE training. **d**. Firing rates of 5 random neurons are plotted for the three different phases of FORCE training (pre-training, during training and post-training). The firing rates before training do not lie in the chaotic regime. They become coherent after entering into the training process. **e**. During training, the outcome trajectories match the Lorenz system. After RLS was turned off, the trajectories deviate. **f**. Chaotic attractors in the three different phase planes post-training. Despite the deviation of the trajectories post-training, the attractors closely resemble the Lorenz system.

### FORCE learning with spiking neural networks of ring topologies

Having shown that FORCE can train ring networks of rate equations, a natural question is whether it can also train ring networks of spiking neurons. In Fig. 3, we compare a spiking ring network with a spiking E-I balanced network. Both networks consist of leaky integrate-and-fire (LIF) neurons and are trained to reproduce the FitzHugh-Nagumo dynamics (see Methods). In the ring network, the neurons were exclusively inhibitory with one neuron inhibiting the next in a ring architecture (Fig. 3a). In the E-I balanced network, excitatory and inhibitory neurons are randomly connected (Fig. 3c). Both networks learned to reproduce the target dynamics (Fig. 3b,d).

**Figure 3:**
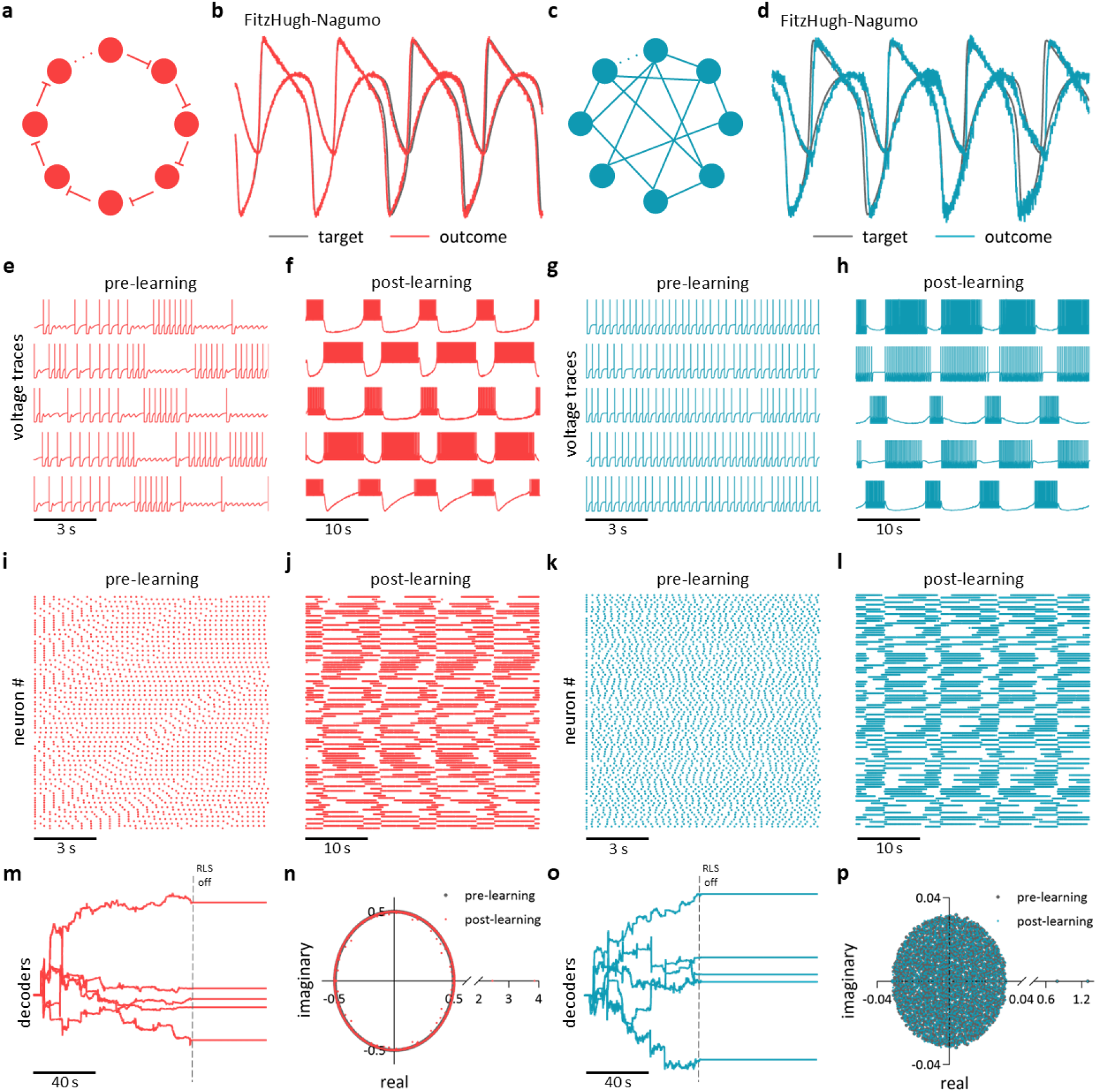
Comparison of the ring spiking neural network and the E-I balanced spiking network. **a**. Schematic illustration of the ring spiking neural network. The network consisted of *N* = 2000 LIF neurons. **b**. The network was trained to reproduce the FitzHugh-Nagumo dynamics. The network outcome is shown in red and the FitzHugh-Nagumo target in grey. **c**. Schematic illustration of the E-I balanced spiking neural network. This network also consisted of *N* = 2000 LIF neurons. **d**. Network outcome (blue) and target dynamics (grey) for the FitzHugh-Nagumo system. **e-f**. Voltage traces pre- and post-training of 5 neurons picked randomly from the ring network. The network was initialized in a non-chaotic, asynchronous regime. Post-training (panel f), the activity becomes coherent. **g-h**. The E-I balanced network initially lies in a chaotic regime, which transforms to coherent during and post-learning (panel h). **i-j**. Raster plot of the spikes pre- and post-learning of the first 100 neurons of the ring LIF network. **k-l**. Corresponding raster plot of the spikes pre- and post-learning of the first 100 neurons of the E-I balanced network. **m**. Decoders of the ring spiking network for the entire duration of the simulation (40s). **n**. Spectrum of the ring network. Eigenvalues pre-learning are shown in grey and post-learning in red. **o**. Decoders of the E-I balanced network. The simulation runs for a total of 40s. **p**. Spectrum of the E-I balanced network. The eigenvalues pre-learning (grey) and post-learning (blue) lie within a circle, with two positive real eigenvalues above 1.

These two networks, despite similar performance, function differently. Prior to learning, spiking activity in the ring network is highly structured (Fig. 3e,i), while activity in the E-I balanced network is irregular (Fig. 3g,k). Following learning, spike trains become regular in both networks (Fig. 3f,j for the ring network and Fig. 3h,l for the E-I balanced network). In both networks, the decoders remain fixed after RLS is turned off (Fig. 3m,o). The eigenvalues of the ring network are located on the shell of a circle (not at the interior) both before and after learning (Fig. 3n), whereas the corresponding spectrum of the balanced network forms a circle in the complex plane (Fig. 3p). For both networks, the leading eigenvalues after learning lie outside the circle-shaped spectra. Once again, ring-like structures with *O*(*N*) parameters in total operate similarly to densely coupled reservoirs with *O*(*N*^2^) parameters.

### Ring spiking neural networks learn low-dimensional dynamics

Next, we sought to determine if spiking rings could robustly learn other tasks. We trained unidirectional ring networks (Fig. 4a) composed of LIF inhibitory neurons to learn multiple dynamical systems tasks, including sinusoidal oscillators, systems with inputs, nested oscillations, and chaotic attractors.

**Figure 4:**
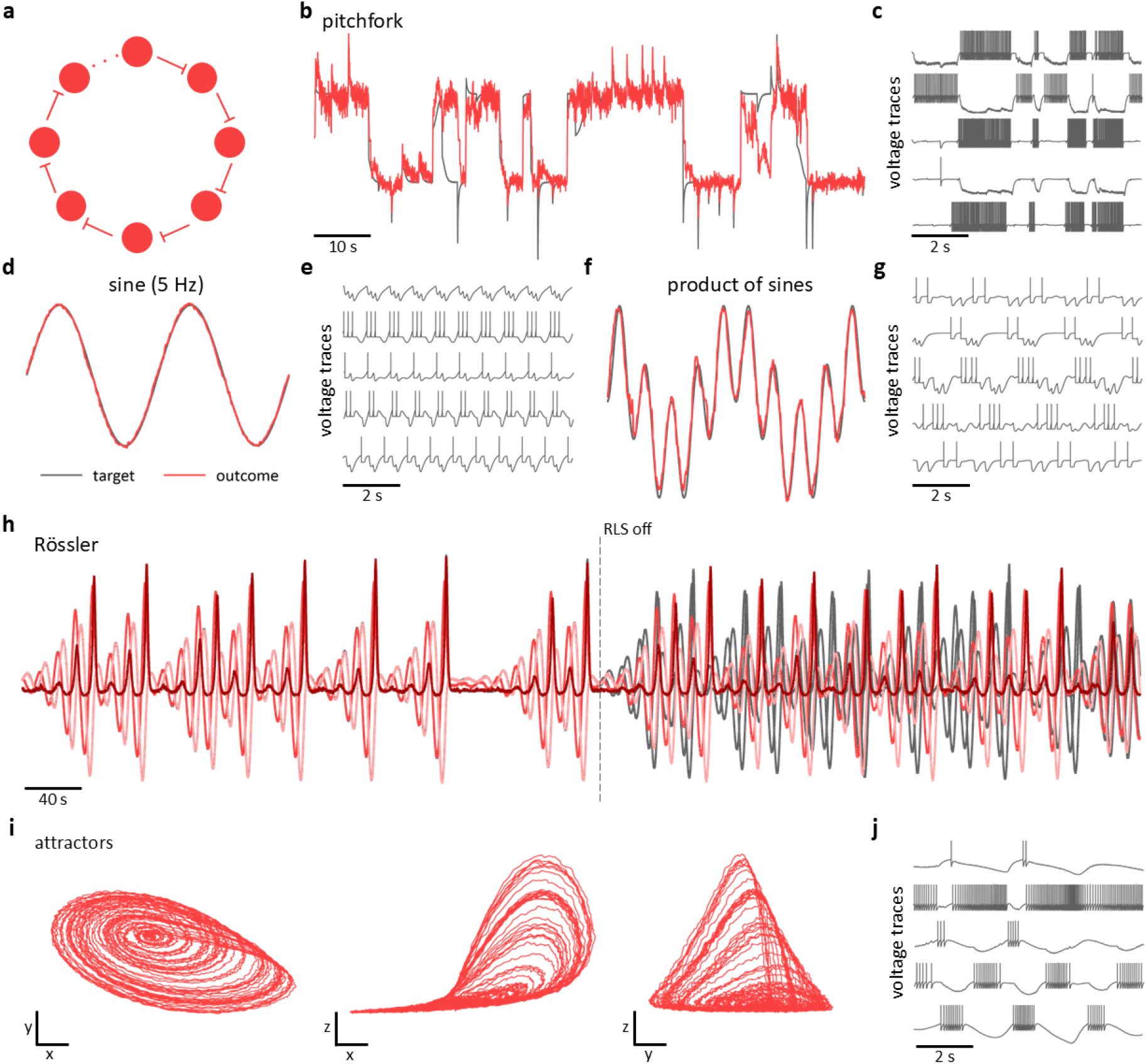
Ring spiking neural networks can train different dynamical systems tasks. **a**. Schematic of a complete ring network. The number of neurons varies from task to task. **b**. The network was trained to reproduce the pitchfork bifurcation. The network consists of a total of *N* = 2000 neurons. Shown is the target and network outcome for 10s post-learning. **c**. Voltage traces for 2s of five randomly picked neurons. **d-e**. In panel **d**, the outcome and target trajectories are shown for the 5Hz sine function. The voltage traces of five random neurons are shown in panel **e** for 2s post-training. **f-g**. Another oscillator was used as a supervisor to train the network. The supervisor is the product of a sinusoid of 4Hz frequency and a sinusoid of 6Hz frequency. Voltage traces of five randomly picked neurons are shown in panel **g. h**. The ring LIF network was trained to reproduce low-dimensional chaotic dynamics. The network outcomes match the Rö ssler trajectories during training but substantially deviate post-training (after RLS is turned off). Chaotic attractors are plotted in three different phase planes, mimicking well the Rö ssler attractors. The displayed attractors are plotted for the time post-training. **j**. Voltage traces post-learning of five neurons randomly picked from the network of *N* = 3000 neurons.

First, we found that FORCE-trained spiking ring networks successfully switch between discrete states and process inputs. A network was trained to reproduce the pitchfork normal form (Fig. 4b), a dynamical system capable of switching between two discrete states with a sufficiently strong input. The network output closely matched the target dynamics qualitative behaviours, with individual neurons also shown to be switching between different states (Fig. 4c).

We additionally trained the networks on periodic oscillatory tasks, including a 5 Hz sinusoid and the product of two sinusoids with fast and slow frequencies (see Methods). Using FORCE, the LIF ring networks successfully learned both the 5 Hz sinusoid (Fig. 4d,e) and the product of sinusoids (Fig. 4f,g).

We further examined the ability of FORCE trained rings to capture temporal patterns by training spiking ring networks to reproduce oscillatory inputs presented repeatedly to the network. As an illustrative example, we took the opening bar of Ode to Joy [14], encoded as a sequence of pulses in a five-dimensional supervisory signal. A ring network of LIF neurons successfully generated this sequence (Supplementary Fig. S2a).

Lastly, we tested the ability of the model to learn more complex dynamics, including low-dimensional chaotic attractors. We considered two examples: the Rössler attractor and the Lorenz attractor (Fig. 4h-j and Supplementary Fig. S2b,c). During the training phase (before RLS is turned off), the readout dynamics of both attractors closely matched the target outputs. After learning (once RLS was deactivated), the outputs diverged from the target trajectories due to the chaotic nature of the attractors (Fig. 4h and Supplementary Fig. S2b). Nevertheless, when the post-learning trajectories are plotted, they closely resemble the corresponding Rössler and Lorenz attractors (Fig. 4i and Supplementary Fig. S2c). For the Rössler attractor, voltage traces from the first five neurons are shown in Fig. 4j.

### FORCE learning in pruned ring networks

We next examined ring networks in more biologically relevant architectures where the rings either have missing connections, are not fully unidirectional, or have some extended non-local connectivity.

First, we considered unidirectional rings with missing connections. A schematic of a unidirectional sparse ring network is shown in Fig. 5a. A network of LIF neurons was trained with FORCE to reproduce multiple target dynamics with 10% of the connections randomly deleted. For the FitzHugh-Nagumo model (Fig. 5b), the post-training activity of the first five neurons is shown in Fig. 5c and is characterized by irregular bursting. Training was achieved using exclusively inhibitory feedforward connections (*Q* < 0). The mean RMSE between the target and the network output remained close to 1 for *Q* ∈ (−5, −0.5), with a minimum value of 0.7 at *Q* = −4 (Fig. 5d). The network was also trained to reproduce the pitchfork normal form (Fig. 5e). Consistent with the FitzHugh-Nagumo case, the mean RMSE remained close to 1 for *Q* ∈ (−5, −0.5), attaining a minimum value of 0.8 at small negative *Q* (specifically, *Q* = −0.5, Fig. 5g). Even with 10% of the existing connections missing, the model could still reproduce the chaotic Rössler attractor (Fig. 5h-k).

**Figure 5:**
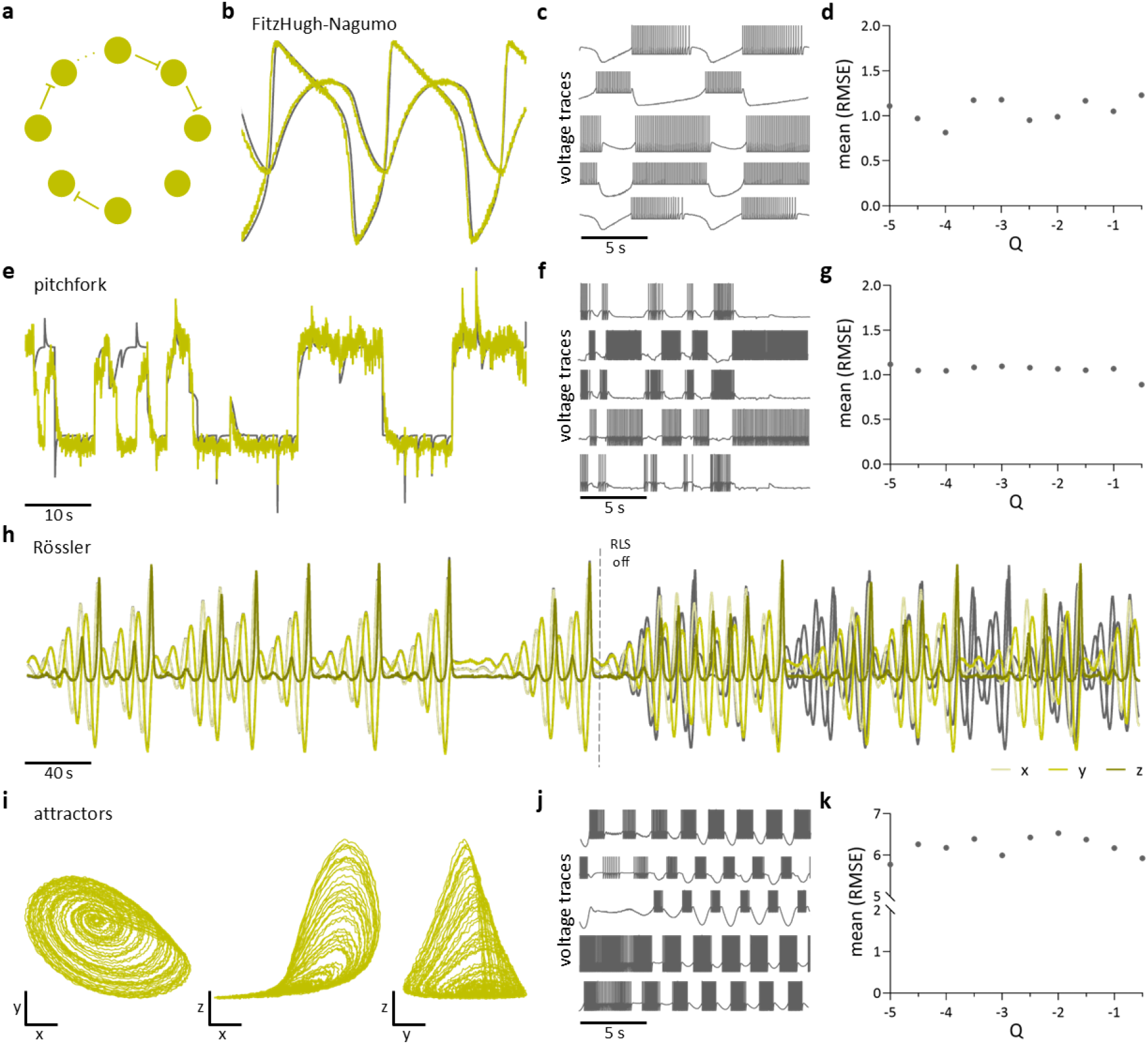
Sparse unidirectional ring networks of spiking neurons effectively learn dynamical systems tasks. **a**. Schematic of a sparse unidirectional ring network. **b**. Trajectories of the target (grey) and the network outcome (yellow) of the FitzHugh-Nagumo system. **c**. Voltage traces of the first five neurons of the network (post-training) for 5 seconds. **d**. Mean value of the RMSE as a function of the ring matrix component *Q*. **e**. The pitchfork normal form target (grey) and the network outcome (yellow). **f**. Voltage traces of the first five neurons showcasing bursting activity post training. **g**. Mean value of the RMSE for a range of *Q* values. **h**. Trajectories of the Rössler system (grey) during and post learning, and corresponding trajectories of the network outcome (yellow shades). **i**. Network attractors of Rössler supervisor post-training. **k**. Mean of RMSE as a function of *Q*.

Next, we considered networks with bidirectional connectivity, in which each neuron is coupled to its immediate neighbours in both directions (Supplementary Fig. S3a). The network was trained to reproduce both the FitzHugh-Nagumo system and the pitchfork normal form. Prior to learning, the coefficient of variation (CoV) of interspike intervals was below 1 for both supervisors, indicating regular or periodic firing (Supplementary Fig. S3b, d). Following learning, CoV values increased above 1, consistent with bursting activity. The RMSE for the FitzHugh-Nagumo task decreased for small negative values of 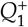 and 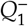 (Supplementary Fig. S3c). Here, the superscripts denote the directionality of the connections, that is, 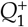 is the coefficient of the forward connections while 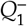 is the corresponding coefficient of backward connections in the bidirectional ring. No significant dependence on either 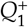 or 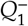 was observed for the pitchfork task (Supplementary Fig. S3e).

Random pruning of connections in the fully connected bidirectional ring yielded a sparse bidirectional ring network (Fig. 6a). Using FORCE, this network was trained to reproduce the FitzHugh-Nagumo dynamics, the pitchfork normal form, and the Rössler attractor (Fig. 6b,e,h,i). Post-learning network activity (Fig. 6c,f,j) exhibited irregular bursting across all three supervisors. Error analysis revealed that, for the

**Figure 6:**
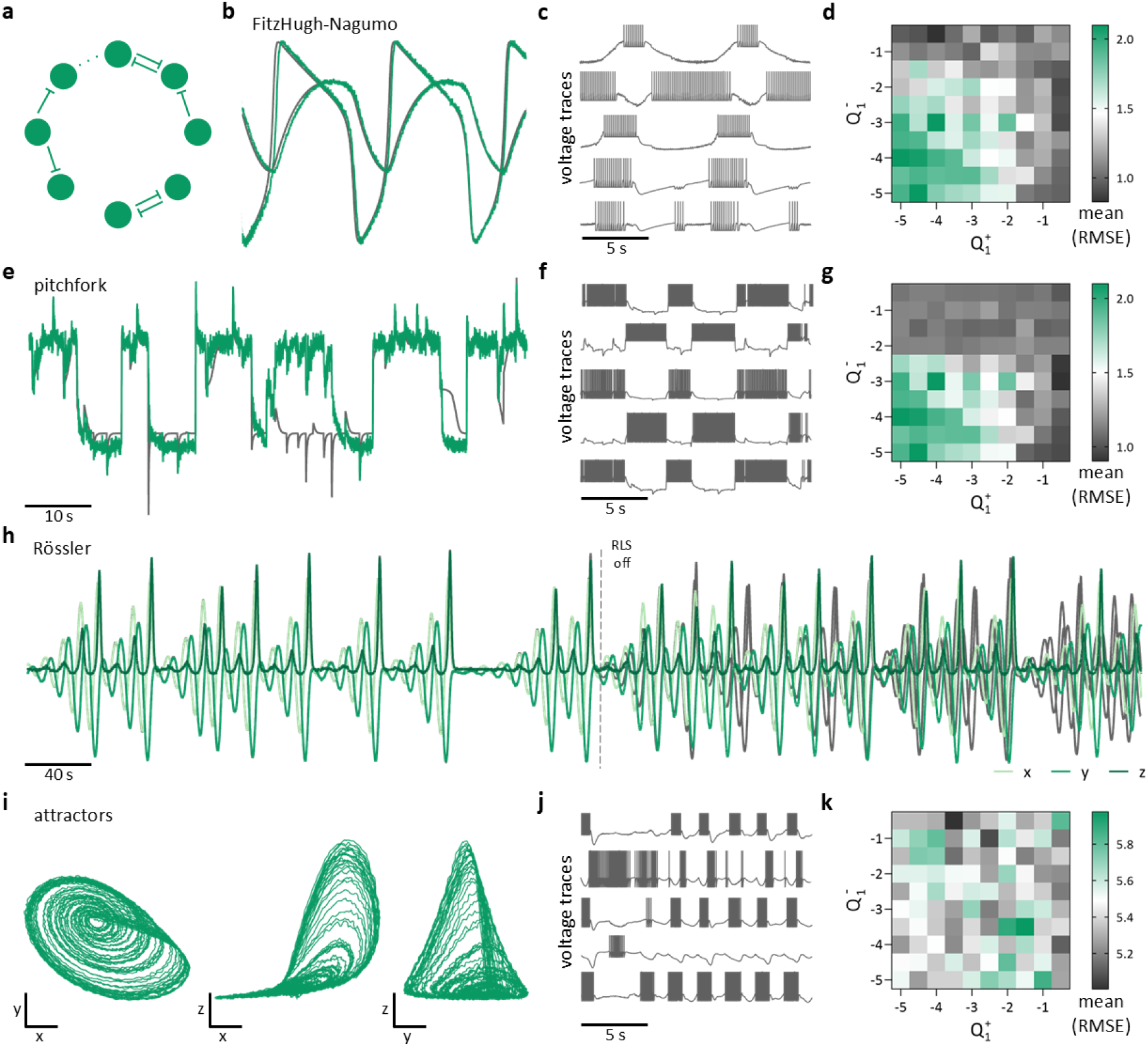
Bidirectional ring spiking neural networks with pruned connections learn different dynamics. **a**. Schematic of a sparse bidirectional ring network. **b**,**e**,**h**. Trajectories of the target (grey) and the network outcome (green) of the FitzHugh-Nagumo (panel **b**), pitchfork (panel **e**) and Rössler system (panel **h**). **c**,**f**,**j**. Voltage traces of the first five neurons (post-training) for 5 seconds. **d**,**g**,**k**. Mean value of RMSE over a grid of points in the 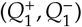 parameter space. **i**. Rössler network attractors post-training.

FitzHugh-Nagumo and pitchfork tasks, the RMSE decreased for small negative values of 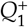 and 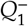 (Fig. 6d,g). In contrast, the RMSE for the Rössler task was larger than for the other two supervisors and with no specific pattern (Fig. 6k).

Finally, we studied bidirectional ring networks with connections extending to both nearest and next-nearest neighbors, as well as a sparse variant of this architecture (Supplementary Fig. S3f, k). All networks could reproduce the FitzHugh-Nagumo dynamics and the pitchfork normal form. As in previous cases, neuronal activity prior to training exhibited regular spiking (CoV < 1), while after training this transitioned to irregular bursting (CoV > 1)(Supplementary Fig. S3g,i,l,n). The RMSE for both supervisors followed trends similar to those observed for the above networks (Supplementary Fig. S3m,o).

Collectively, these results show that ring structures can learn dynamics as effectively as densely coupled balanced recurrent neural networks, utilizing techniques from reservoir computing to learn a fixed decoder. This learning is also robust to imperfections and asymmetries in the rings.

## Discussion

In this work, we studied ring networks of rate equations and spiking neural networks, demonstrating that rings could effectively learn dynamics, in many cases as well as densely coupled recurrent neural networks with reservoir computing. The spatio-temporal activity patterns produced by rings can be used to learn and reproduce a broad range of dynamics including chaotic attractors, oscillatory attractors, and multi-stable switches.

More specifically, in rate networks, our results show that the success of FORCE training is not uniquely tied to the presence of dense and balanced connectivity that produces a chaotic asynchronous state. While the asynchronous state, as in the original formulation of FORCE, can provide a rich substrate for learning, our findings indicate that structured spatio-temporal activity patterns produced by rings and a shell of eigenvalues can also support the learning of dynamical systems trajectories. Extending these results to spiking neural networks, we found that FORCE training is comparably effective in ring architectures composed of LIF neurons, despite their markedly different microscopic dynamics compared with randomly connected E-I balanced networks. Notably, the ring spiking neural networks successfully reproduce the target dynamics even though the pre-learning activity is highly structured rather than chaotic. Following learning, both the ring and E-I balanced networks converge to coherent spiking patterns, indicating that FORCE training can stabilize task-relevant activity across distinct architectural and dynamical regimes. Hence, we may conclude that simple spiking ring networks may serve as expressive and flexible substrates for learning dynamical systems. These results were also robust when considering rings with missing connections, and rings with random or bidirectional connections. Together, these results highlight the capacity of sparse but structured networks to learn and sustain rich dynamics, with potential relevance for ring attractor representations underlying head-direction coding in systems such as the Drosophila central brain [30, 31] and the zebrafish heading-direction circuit [29].

While the present results show that FORCE training performs well in networks composed of LIF neurons, future work should assess how the choice of neuronal model influences learning performance. Previous work has shown that networks based on the Izhikevich neuron model can achieve higher accuracy when learning dynamical systems [14], owing in part to the presence of spike-frequency adaptation variables that introduce slower intrinsic timescales. Such additional timescales may enhance the expressive capacity of recurrent networks by enriching their reservoir dynamics. More generally, other biologically motivated mechanisms that operate on longer timescales, such as synaptic depression, may similarly increase the ability of ring spiking networks to support supervised learning [49–51].

Beyond its performance on dynamical systems, our approach may provide a useful tool for modeling and interpreting experimentally observed neural dynamics. Recent work has revealed ring attractor networks of heading direction in zebrafish [29], including topographically organized neural activity that evolves smoothly over time and remains stable over extended periods. Such experimental observations highlight the relevance of simple recurrent architectures with inhibitory connectivity for representing low-dimensional variables in the brain. In this context, the ring networks studied here could be used to fit and reproduce experimental data. Applying this approach to neural recordings may therefore offer a principled way to link circuit architecture, learned dynamics, and behavior, and to test hypotheses about how ring attractor computations are implemented in biological networks.

Related experimental work in Drosophila has provided compelling evidence for ring-attractor circuits that rely on local excitation combined with global inhibition to sustain and update heading-direction representations [30, 31]. In these systems, population activity forms a stable, bump-like pattern whose position can be continuously shifted by sensory or motor signals. The computational framework presented here could be used to capture such dynamics, and could be extended to incorporate structured excitation and inhibition consistent with experimentally identified circuit motifs. Such an approach may help bridge theoretical models of ring attractors with experimentally observed circuit mechanisms in the fly brain.

Finally, we remark that our work has relevance for neuromorphic instantiations of reservoir computing. In lieu of potentially requiring *N*^2^ fixed connection weights which are binary (inhibitory/excitatory), ring reservoirs allow for robust computation with only *O*(*N*) unary (inhibitory) weights. This can considerably simplify complex circuit designs while still producing rich enough dynamics to support learning.

## Methods

### Networks of rate equations

We consider the following system of rate equations:

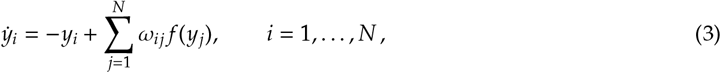

where *y*_*i*_ is interpreted as the membrane potential of the *i*-th cell, for a total of *N* cells. The synaptic weight matrix *ω*_*ij*_ couples the output of presynaptic neuron *j* to the input of postsynaptic neuron *i*. The function *f* (*y*), denoting the firing rates, is given by a sigmoid function of the form:

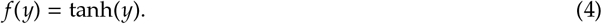

The synaptic weight matrix *ω* can be expressed as the sum of static and learned components:

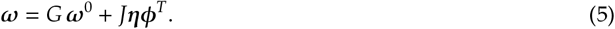

The static component is defined by the static weight matrix *ω*^0^, which is a random matrix scaled by the gain parameter *G*, while the learned component is given by the product of a static random encoder *η* and a learned decoder *ϕ*.

The weight matrix *ω*^0^ is drawn from the normal distribution with mean and variance given respectively by

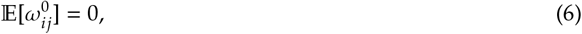

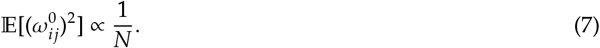

The first equation implies that excitatory weights 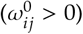 approximately balance the inhibitory weights 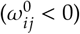, while the second equation forces the neurons to be strongly coupled, which leads to the asynchronous, irregular firing displayed by both spiking and rate networks. The decoders *ϕ* are iteratively learned with the RLS technique, described in detail below. The encoders *η* are drawn from a uniform distribution over [−1, 1], specifying each neuron’s encoding preference across the phase space of the target dynamics.

#### Ring topologies

The recurrent neural network topologies considered in this study have a ring structure, with neurons arranged in a circular configuration. Several variants are examined. The matrix 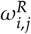 for a unidirectional ring with one-way connectivity forming a closed loop is defined by

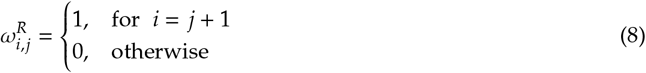

with 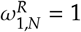. For a bidirectional ring in which each neuron connects to its immediate neighbors in both directions, the matrix 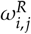 takes the form

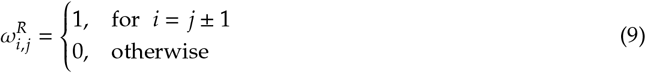

with 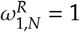 and 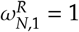. While for a bidirectional ring with connections extending to both nearest and next-nearest neighbors, such that each neuron is linked to its two immediate neighbors as well as to the two neurons adjacent to them, the matrix becomes

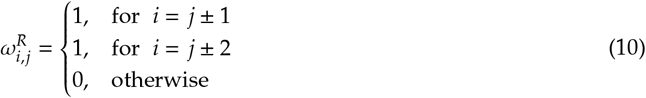

with 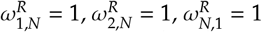 and 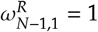.

We further considered a sparse version of each of the above three ring network topologies. Existing connections were removed uniformly at random with probability 0.1.

#### Networks of LIF neurons

The spiking networks consist of leaky integrate-and-fire (LIF) neurons of the form

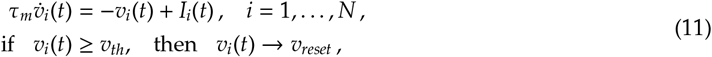

where *v*_*i*_ is the membrane potential of the *i*-th neuron. When the membrane potential *v*_*i*_ reaches the threshold value *v*_*th*_, the neuron emits a spike, after which *v*_*i*_ is reset to *v*_*reset*_ and held there for a refractory period *τ*_*ref*_. The membrane time constant *τ*_*m*_ plays a central role in shaping the neuron’s response to inputs. The parameter values are listed in Table 1.

**Table 1.** List of parameter values of the LIF neuron model.

| Symbol | Value |
| --- | --- |
| $\tau_m$ | 0.01 s |
| $\tau_{ref}$ | 0.03 s |
| $v_{reset}$ | -65 mV |
| $v_{th}$ | -40 mV |
| $I_{bias}$ | -39.5 pA |
| $\tau_R$ | 0.002 s |
| $\tau_D$ | 0.03 s |

The spikes are filtered using the double exponential synaptic filter:

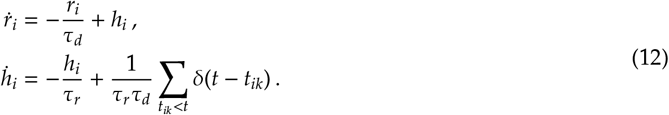

The parameters *τ*_*r*_ and *τ*_*d*_ represent the synaptic rise time and the synaptic decay time, respectively, and *δ*(·) is the Dirac delta function.

The current *I*_*i*_ is defined as *I*_*i*_(*t*) = *I*_*bias*_ + *s*_*i*_(*t*). Here, *I*_*bias*_ is a constant background current set at a value slightly above the rheobase to ensure excitability, and *s*_*i*_(*t*) are the synaptic input currents given by

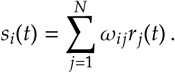

The synaptic connectivity matrix *ω*_*ij*_ determines the strength of post-synaptic currents transmitted from neuron *j* to neuron *i*.

#### FORCE training

The goal of FORCE training is to align the network output 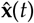 with the trajectory of the target dynamics **x**(*t*). This is achieved by minimizing the squared error between the two signals while constraining the magnitude of the decoder weights. The loss function is given by

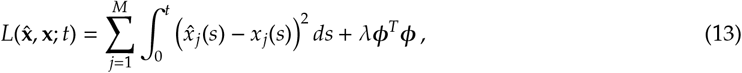

where *λ* is a regularization coefficient. To update the decoder *ϕ*, FORCE uses the Recursive Least Squares (RLS) algorithm, which reduces the prediction error

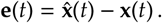

At discrete time steps of size Δ*t*, the updates are computed as

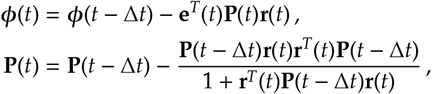

where **P**(*t*) serves as a discrete approximation to the inverse of the correlation matrix with regularization *λ*:

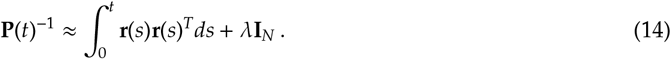

In Eq. (14), **I**_*N*_ is an *N*-dimensional identity matrix. The decoders and correlation matrix are initialized as

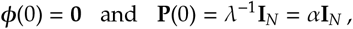

respectively. The parameter *α* is a learning rate that controls how fast FORCE adapts during training.

#### Teaching signals

To train the ring networks of spiking LIF neurons, we considered multiple dynamical systems tasks given in detail below.

##### Sine wave

A simple sine wave of 5Hz,

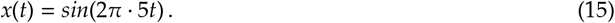

##### Product of sine waves

We considered the product of two sine waves as a supervisor:

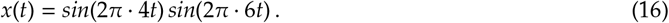

##### FitzHugh-Nagumo

The FitzHugh-Nagumo model is a simplified model of neuronal excitability. The equations are given by:

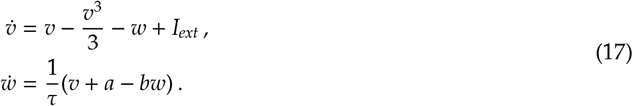

The variables *v* and *w* describe the membrane potential and recovery variable, respectively. The parameters *a* and *b* shape the nullclines and recovery behaviour, *τ* is the time scale separation (making recovery slower than excitation), and *I*_*ext*_ is the external input current. The parameter values used are *a* = 0.2, *b* = 0.8, *τ* = 10, and *I*_*ext*_ = 0.5.

##### Ode to Joy

A 5-dimensional teaching signal consisting of a sequence of pulses that correspond to the notes in the first bar of the song [14]. The positive component of a sine wave with a frequency of 2Hz was used for quarter notes and with 1Hz for half notes. Each wave corresponds to the presence of a note and they can also be used as the amplitude of the harmonics corresponding to each note to generate an audio-signal from the network.

##### Pitchfork

The pitchfork normal form with an external random forcing *p*(*t*) was used to generate the pitchfork teaching signal:

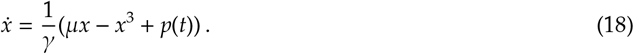

The time-scale parameter *γ* changes how fast trajectories move toward equilibria or away from unstable states, and how they react to the random forcing. The parameter *µ* is the bifurcation parameter. Without forcing (i.e., for *p*(*t*) = 0), and for *µ* > 0 there are three equilibrium points: 0 (unstable) and ± 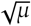 (stable). In this bistable regime, adding the perturbations *p*(*t*) can push the trajectory back and forth between the two wells, that is, around the points ± 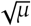. We fixed the parameter values to *γ* = 0.06 and *µ* = 1. For the perturbations *p*(*t*), we generated a train of random square pulses. The intervals between pulses were chosen uniformly from [0.1, 0.3]. Each pulse has a short duration sampled uniformly from [0.007, 0.01] and an amplitude that is drawn from a normal distribution with mean 0 and variance 2.

##### Rössler system

The Rössler system consists of the following three coupled nonlinear differential equations:

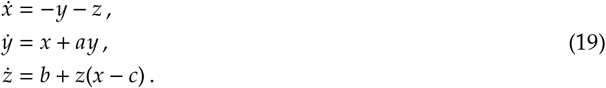

The parameter values used are *a* = 0.432, *b* = 2, and *c* = 4.

##### Lorenz system

The equations of the Lorenz system are given by:

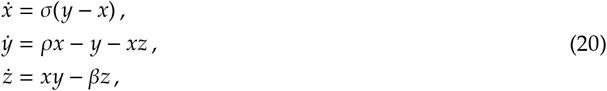

with parameter values *σ* = 10, *ρ* = 28, and 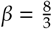.

#### Simulation details for Figure 1 & 2

A network governed by rate equations was trained to reproduce the FitzHugh-Nagumo dynamics and the Lorenz chaotic attractor. For the simulation corresponding to Fig. 1, the network size was *N* = 1000 with Δ*t* = 2 ms and a total duration of 10 s. Learning was on from 1 s to 5 s.

For the simulation corresponding to Fig. 2, the network size was *N* = 2000 with Δ*t* = 2 ms and a total duration of 17 s. The weight matrix *ω* was parameterized by *G* = 0 and *Q* = 1.5, forming a ring reservoir. Learning occurred from 1 s to 8.5 s, followed by reproduction of the target signal until the end of the simulation. In both cases, the learning rate was *α* = 1 ms.

#### Simulation details for Figure 3

The ring spiking neural network consisted of *N* = 2000 neurons. The total simulation time was 50 s with Δ*t* = 0.1 ms. With *G* = 0, *Q* = −0.5 and *J* = 60, the network formed a purely inhibitory ring. Training was performed for 20 s, from 10 s to 30 s, followed by reproduction of the FitzHugh-Nagumo dynamics from 30 s to 50 s.

An E-I balanced network with the same size (*N* = 2000) was also tested. The training interval was identical. The difference lay in the weight matrix *ω*, with parameters *G* = 0.02, *Q* = 20 and *J* = 20, and network sparsity *p* = 0.1. For both networks, the numerical integration time step was 0.05 ms, and *α* = 0.001 ms.

#### Simulation details for Figure 4

Networks of LIF neurons were trained to reproduce several supervisors. For the pitchfork supervisor, the network comprised *N* = 2000 inhibitory neurons with *G* = 0, *Q* = −21, *J* = 20 and Δ*t* = 0.05 ms. Training was performed from 1 s to 30 s, followed by reproduction of the temporal dynamics from 30 s to 50 s.

For the 5 Hz oscillator, the network comprised *N* = 1000 inhibitory neurons with *G* = 0, *Q* = −2, *J* = 60 and Δ*t* = 0.1 ms. Training occurred from 5 s to 15 s, after which the oscillator was reproduced from 15 s to 30 s. For the second oscillator (defined as the product of two sinusoids), the network contained *N* = 2000 inhibitory neurons with connectivity parameters *G* = 0, *Q* = −2, *J* = 30 and Δ*t* = 0.2 ms. The total simulation time was 40 s, with training from 1 s to 30 s and reproduction during the remaining 10 s.

For the Rössler attractor, the network comprised *N* = 3000 inhibitory neurons with *G* = 0, *Q* = −30, *J* = 10 and Δ*t* = 0.2 ms. The total simulation time was 80 s, including a training phase from 1 s to 40 s, followed by reproduction of the Rössler trajectories for the remaining duration.

For all supervisors, the numerical integration time step was 0.1 ms and *α* = 0.001 ms.

#### Simulation details for Figure 5

A sparse unidirectional ring network of LIF neurons was trained to reproduce several dynamical systems. For the FitzHugh-Nagumo system, we used *N* = 2000, *Q* = −0.2, *J* = 60 and Δ*t* = 0.3 ms. The network was trained from 1 s to 20 s and subsequently reproduced the FitzHugh-Nagumo trajectories from 20 s to 40 s. For the pitchfork dynamics, we used *N* = 2000, *Q* = −0.2, *J* = 20 and Δ*t* = 0.05 ms. The network was trained from 1 s to 30 s and reproduced the dynamics for 20 s. For the Rössler attractor, the network size was *N* = 3000, with *Q* = −0.5, *J* = 10 and Δ*t* = 0.2 ms. The network reproduced Rössler-like attractors for 40 s after 39 s of training. For all supervisors, the integration time step was 0.1 ms, *α* = 0.001 ms, and *G* = 0.

#### Simulation details for Figure 6

As in Fig. 5, a sparse bidirectional ring network of LIF neurons was used to generate the results shown in Fig. 6 for three dynamical systems: the FitzHugh-Nagumo system 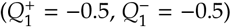, the pitchfork dynamics 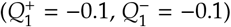, and the Rössler attractor 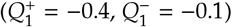. The network size *N*, gain parameters *Q* and *J*, and simulation time step Δ*t* were identical to those in Fig. 5 for all supervisors. The numerical integration time step was 0.1 ms, with *α* = 0.001 ms and *G* = 0.

## Data Availability

No data was generated by this study.

## Code Availability

The authors will make the code available upon publication.

## Acknowledgments

We would like to thank the Hotchkiss Brain Institute and the Cumming Medical Research Fund for supporting this work.

## Author Contributions Statement

AT wrote the manuscript and performed mathematical analysis and numerical simulations. WN and AT conceived of the study. WN edited the manuscript and supervised the project.

## Declaration of Interests

WN is the CSO of Synaptrain Technologies Inc. Synaptrain Technologies Inc. did not influence the design and results of this manuscript and did not fund this research.

## Supplementary Figures

**Figure S1:**
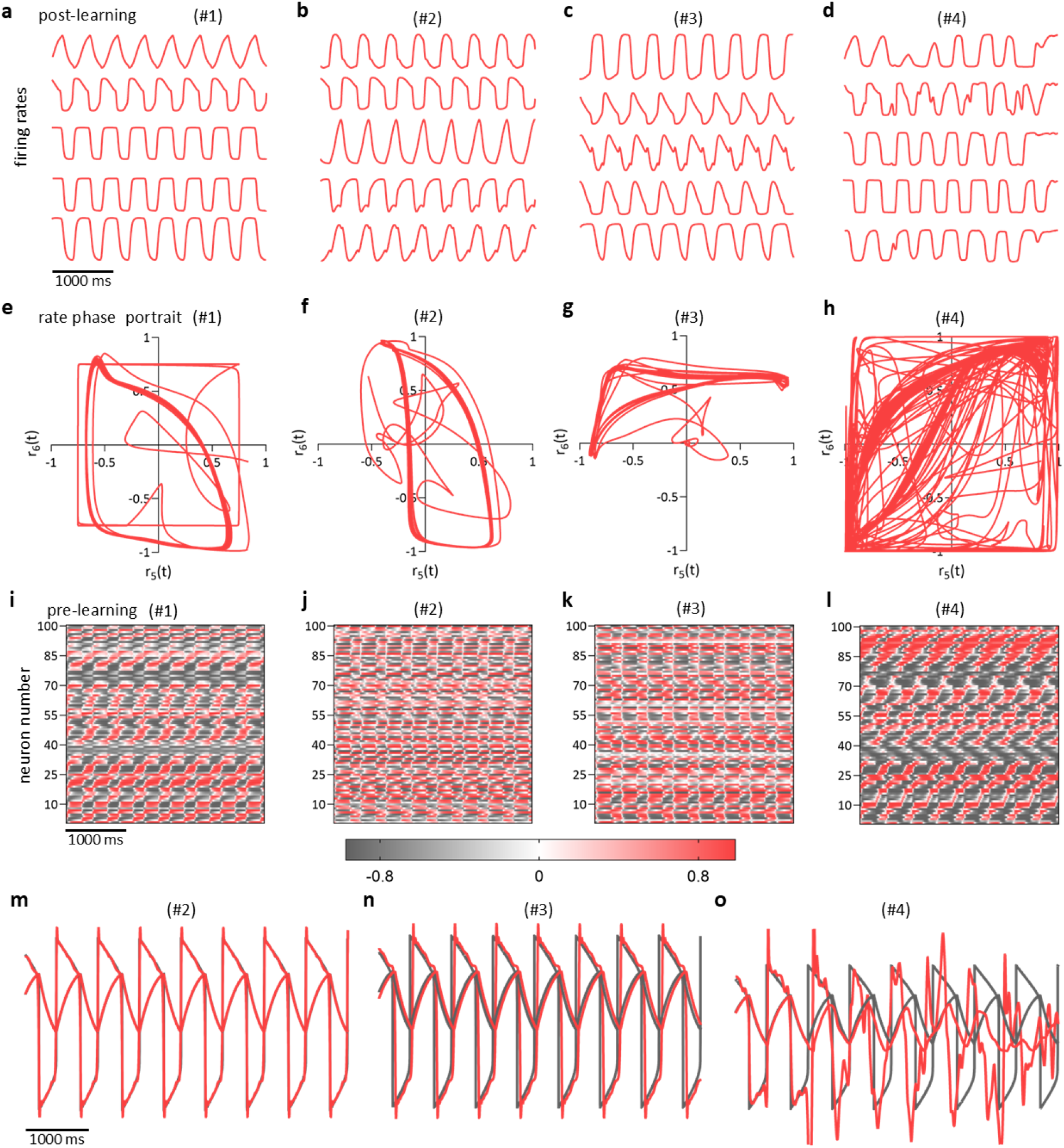
**a-d**. Firing rates for 1000 ms post learning, for the four distinct points picked in Fig. 1. **e-h**. Rate phase portraits for neurons 5 and 6. **i-l**. Patterns of firing rates for 1000 ms before training starts. **m-o**. Target and network outcome trajectories post-training for points #2, #3 and #4.

**Figure S2:**
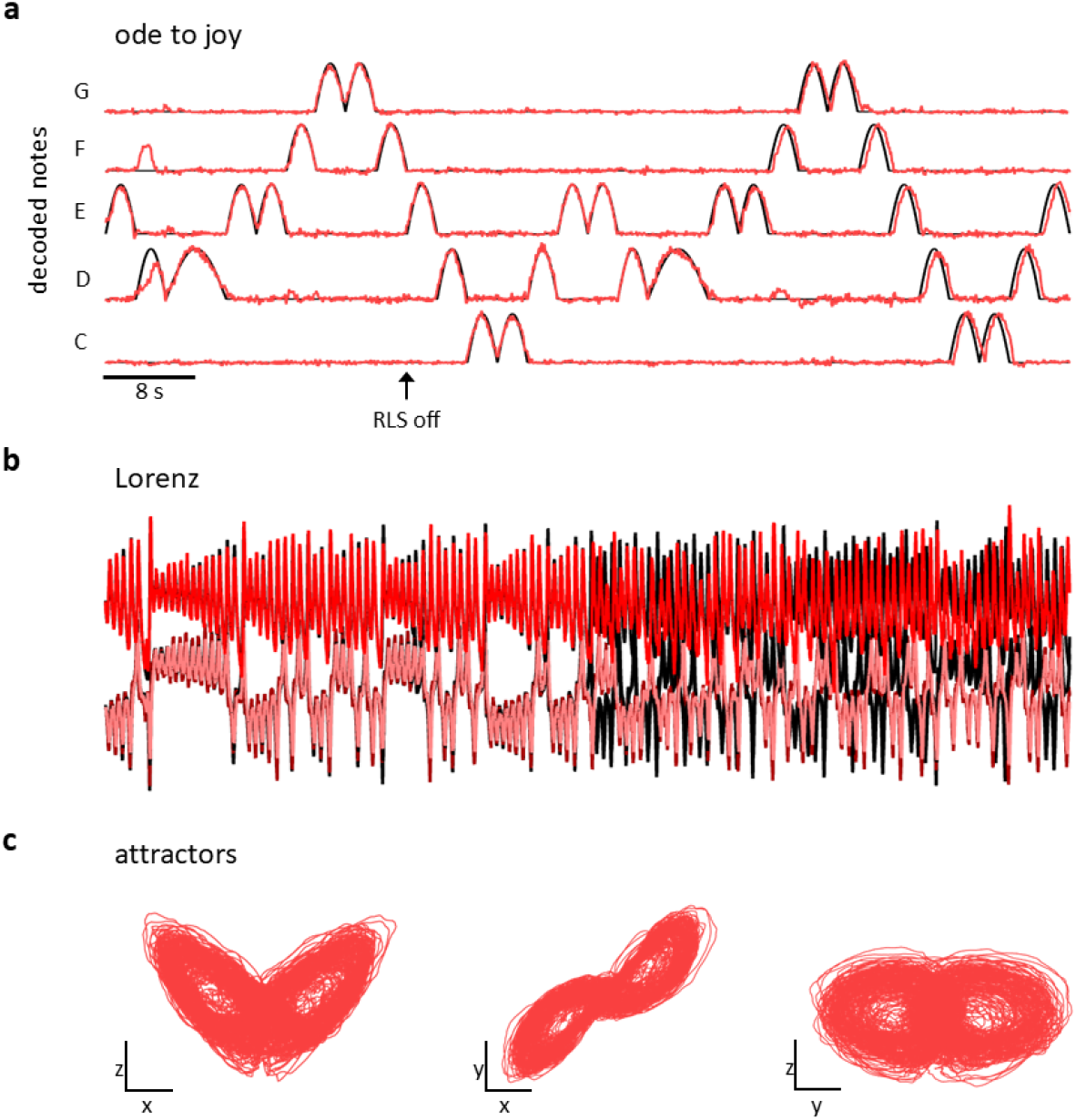
**a**. Teaching signal of five notes of the song Ode to Joy. The network was initialized in a non-chaotic regime and trained to reproduce the signal. Parameter values used for Ode to Joy: *N* = 2000, Δ*t* = 0.5 ms, *G* = 0, *Q* = −0.8. The total simulation time was 300 s, of which 259 s were used for training. **b**. The Lorenz chaotic system used as a teaching signal. Post-training the trajectories diverged due to chaos. **c**. Network attractors post-training. Parameter values used for Lorenz attractor: *N* = 5000, Δ*t* = 0.2 ms, *G* = 0, *Q* = −0.8. The total simulation time was 50 s, of which 24 s were used for training.

**Figure S3:**
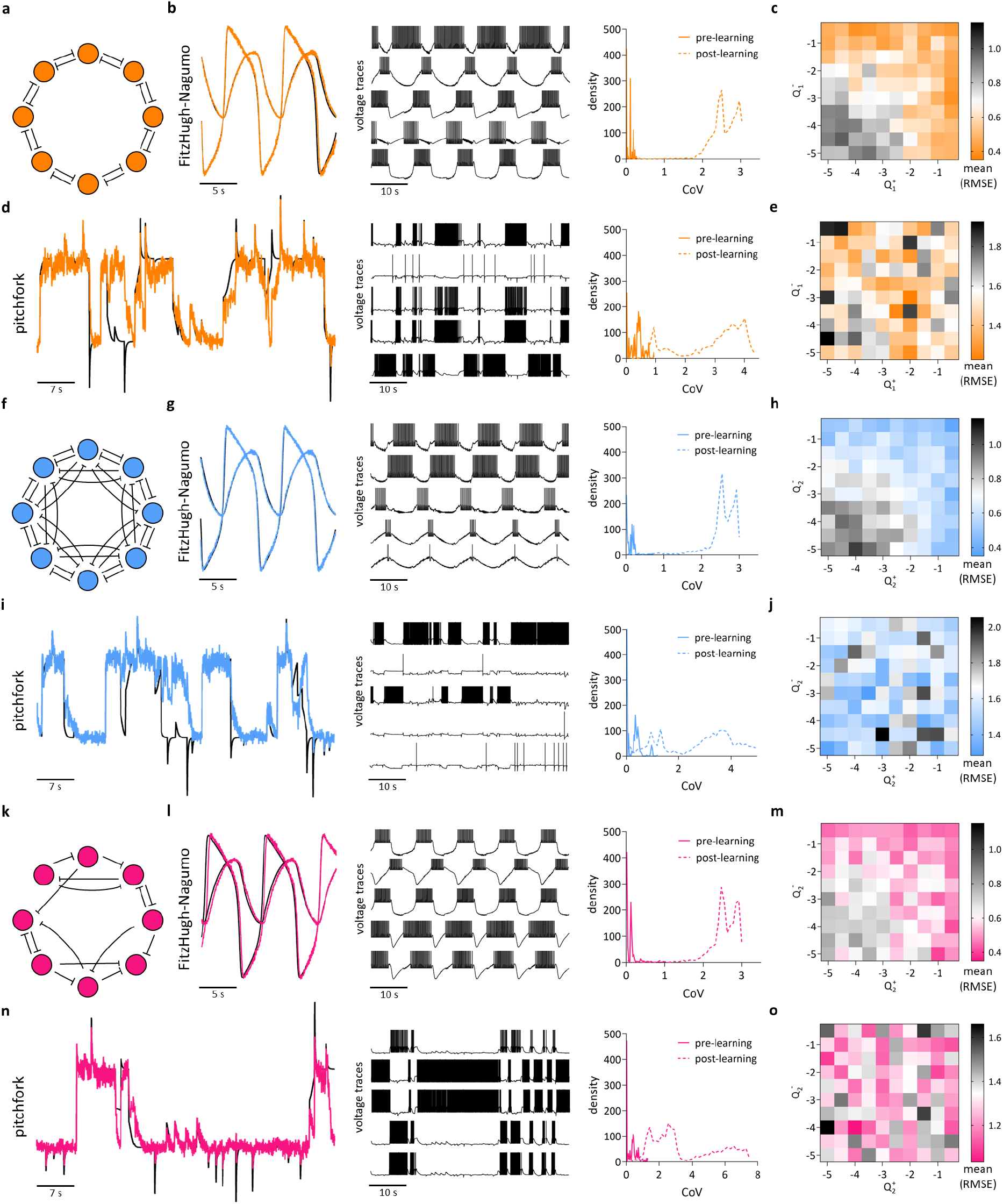
**a**. Schematic of a bidirectional ring network. **b**. Trajectories and voltage traces post training of the FitzHugh-Nagumo system. The third panel in **b** is the density of the coefficient of variation (CoV) pre- and post-learning. Since CoV < 1, the activity pre-learning is regular or periodic. However, it switches to bursting post learning since CoV becomes greater than 1. **c**. Heatmap of the mean RMSE over a grid of points in the 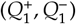 parameter space. **d**. Pitchfork normal form: teaching signal, voltage traces and CoV densities. **e**. Heatmap of the mean RMSE for the pitchfork supervisor. **f**. Schematic of a bidirectional ring with nearest and next-nearest connections. **g**. Trajectories, voltage traces and CoV density for the FitzHugh-Nagumo supervisor. **h**. Heatmap of the mean RMSE for the FitzHugh-Nagumo supervisor. **i**. Teaching signal, voltage traces and CoV density for the pitchfork supervisor. **j**. Heatmap of the mean RMSE for the pitchfork supervisor. **k**. Sparse version of the ring network of panel **f. l**. Trajectories, voltage traces and CoV density for the FitzHugh-Nagumo supervisor. **m**. Heatmap of the mean RMSE for the FitzHugh-Nagumo supervisor. **n**. Teaching signal, voltage traces and CoV density for the pitchfork supervisor. **o**. Heatmap of the mean RMSE for the pitchfork supervisor.

